# Zero-Shot Metabolite Prediction from Gene Expression via Physics-Informed Graph Neural Networks

**DOI:** 10.64898/2026.06.29.735073

**Authors:** Claudio Novella-Rausell, Ton Rabelink, Ahmed Mahfouz

**Affiliations:** Department of Human Genetics, Leiden University Medical Centre, 2333 ZA Leiden, the Netherlands; Department of Internal Medicine (Nephrology) & Einthoven Laboratory of Vascular and Regenerative Medicine, Leiden University Medical Center, Leiden, The Netherlands; The Novo Nordisk Foundation Center for Stem Cell Medicine (reNEW), Leiden University Medical Center, Leiden, The Netherlands; Leiden Computational Biology Center, Leiden University Medical Center, Leiden, the Netherlands; Delft Bioinformatics Lab, Delft University of Technology, Delft, the Netherlands

## Abstract

Predicting metabolite concentrations from gene expression is instrumental for linking regulatory programs to metabolic phenotypes. Prior approaches rely on static enzyme–metabolite mappings and often omit data-driven learning or biochemical constraints, limiting their ability to generalize to new metabolites. We present GAZE (Graph Attention for Zero-shot metabolite Estimation), a physics-informed graph neural network that integrates enzyme expression, Enzyme Commission functional embeddings, and ChemBERTa metabolite descriptors within a unified metabolic graph (5,414 nodes, 16,307 edges). A Metabolite-Conditioned Reader uses each metabolite’s SMILES embedding to query learned pathway representations, enabling zero-shot prediction with no metabolite-specific parameters. We evaluate GAZE in three scenarios: (i) standard cross-validation on the Cancer Atlas of Metabolic Profiles (18,044 genes, 180 metabolites, 867 cell lines), achieving R^2^ = 0.816; (ii) leave-one-metabolite-out (LOMO) zero-shot evaluation across 50 held-out metabolites, where the physics-informed variant halves the median R^2^ deficit relative to a standard GNN baseline (−0.34 vs. −0.72), with 30% of unseen metabolites achieving positive R^2^; and (iii) external validation on an independent clear cell renal cell carcinoma tissue cohort (220 samples), where GAZE achieves median Spearman *ρ* = 0.330 across 214 metabolites without fine-tuning. GAZE outperforms scCellFie, MEBOCOST, and UnitedMet across all evaluation settings.

## 1 Introduction

The metabolome represents the ultimate readout of cellular phenotype, integrating the output of genomic instructions with environmental constraints [1]. While single-cell transcriptomics has revolutionized our ability to map cellular heterogeneity [2], characterizing the metabolome at a similar resolution remains analytically challenging. Metabolomics technologies, although powerful, are often destructive, expensive, and limited in coverage compared to the genome-wide scope of RNA sequencing [3]. This asymmetry has motivated the development of computational methods aimed at inferring metabolic states directly from widely available gene expression data. Bridging this gap is essential for unlocking the metabolic information latent in transcriptomic atlases and for understanding disease mechanisms where metabolic reprogramming plays a central role.

Current computational approaches generally fall into two categories: supervised imputation and knowledge-based inference. Supervised methods, such as LASSO regression [4] and the Bayesian framework UnitedMet [5], treat gene expression as a statistical predictor of metabolite abundance. These models have demonstrated success in identifying metabolic phenotypes, such as oxidative shifts in renal cell carcinoma [5], by learning covariance structures from paired RNA-metabolite datasets. However, they operate under a closed-world assumption: they can only predict metabolites explicitly present in the training set. When a metabolite is entirely absent from training, no learned representation exists for the target, and generalization to novel or unmeasured compounds is not possible without auxiliary information such as chemical structure or network context [6].

In contrast, knowledge-based methods treat gene expression as a direct proxy of metabolic potential. Tools like scCellFie [7] aggregate enzymes into “metabolic tasks”—predefined sets of reactions representing a specific metabolic function (e.g., glutamine degradation)—to score pathway activity, while MEBOCOST [8] infers metabolite-mediated cell-cell communication by examining the co-expression of biosynthetic enzymes and sensor receptors. Although valuable for generating qualitative hypotheses, these methods rely on the strong assumption that the abundance of transcripts is linearly mapped to the metabolic flux, a premise often violated by post-transcriptional regulation and reaction kinetics [3]. Furthermore, these proxy-based frameworks are rarely validated against directly measured metabolomics, relying instead on biological plausibility or downstream functional phenotypes [7, 8].

Both approaches share a fundamental limitation: the inability to generalize predictions to unseen metabolites in a rigorous, chemically grounded manner. To predict the abundance of a metabolite not present in the training data, a model must learn to recognize the underlying chemical and topological rules that govern metabolism rather than simply memorizing gene-metabolite correlations. Graph Neural Networks (GNNs) have successfully addressed similar zero-shot problems in drug response prediction [9, 10] by encoding molecular structures. However, applying this to endogenous metabolism requires integrating chemical representation learning with the physical constraints of metabolic networks [11].

Here, we investigate the architectural determinants of zero-shot generalization in metabolite prediction. We developed a modular Graph Neural Network (GNN) framework to systematically evaluate how enzyme functional embeddings and molecular structural representations can be integrated to predict concentrations of unseen metabolites. We assess the specific contributions of novel architectural components, including cross-modal attention for priming pathway context and a Metabolite-Conditioned Reader (MCR) designed to query the network based on chemical structure. Furthermore, we explore the integration of physics-informed loss functions that enforce mass balance and flux constraints, testing the hypothesis that biochemical consistency is a prerequisite for generalization. Our results disentangle the impact of these design choices, demonstrating that chemically-aware, physics-constrained architectures can match dedicated per-metabolite baselines while enabling zero-shot prediction of the unmeasured metabolome.

## 2 Materials and Methods

### 2.1 Problem formulation

We formalize metabolite prediction as a node regression task on a heterogeneous directed graph *G* = (*V, ε*), where *V* = *V*_*r*_ ∪ *V*_*m*_ comprises reaction nodes *V*_*r*_ and metabolite nodes *V*_*m*_ connected exclusively by cross-type edges (reaction ↔ metabolite). Edges ℰ encode substrate–product relationships annotated with stoichiometric coefficients. Given a gene expression profile **g** ∈ ℝ^*G*^ for a biological sample, the model predicts the concentration vector 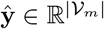 for all metabolites simultaneously, including those with no observed training data.

### 2.2 Data acquisition and preprocessing

We utilized the Cancer Atlas of Metabolic Profiles (CAMP) [12], which pairs untargeted metabolomics (Metabolon LC-MS/MS and GC-MS) with transcriptomic profiles (18,044 genes after filtering to protein-coding genes with nonzero variance) across cancer cell lines. Gene symbols were mapped to Ensembl identifiers to align with the metabolic network gene rules, yielding 988 cell lines with matched expression and metabolomics profiles.

The raw metabolomics data contained 2,504 named biochemicals. We retained the 1,152 metabolites annotated with at least one database identifier (HMDB [13] or KEGG [14]) from the annotation sheet. Samples containing any negative metabolite values (121 cell lines) were removed, leaving 867 cell lines for downstream analysis. Metabolite concentrations were used as provided by Metabolon (median-scaled, log-transformed intensity values). Of the 1,152 annotated metabolites, 380 could be mapped to metabolite nodes in the metabolic network graph via their HMDB or KEGG identifiers; the remaining unmapped metabolites were excluded from the prediction target. Per-sample metabolite measurements that were missing (NaN) were masked during loss computation, so each cell line contributed to the loss only for its non-missing metabolites. On average, approximately 180 of the 380 graph-mapped metabolites were measured (non-missing) per cell line. The 867 cell lines were partitioned into 70/15/15% (train/validation/test) splits via seed-controlled random shuffling. For the modality ablation study, 5-fold cross-validation was used instead.

To evaluate cross-domain generalization, we used two sub-cohorts (RC18, *n* = 144; RC20, *n* = 76) from the UnitedMet clear cell renal cell carcinoma (ccRCC) cohort [5], totalling 220 tumour tissue samples with paired bulk RNA-seq and mass spectrometry metabolomics. Gene symbols were mapped to the same Ensembl identifiers used in the CAMP graph, and metabolite names were matched to graph nodes via HMDB and KEGG identifiers. Raw metabolomics intensities were log_2_(1 + *x*) transformed to approximate the CAMP concentration scale. No gene-expression normalization or domain-adaptation procedure was applied. The model received external expression profiles through the same input pipeline used during training. Of the 685 metabolites in the ccRCC panel, 230 could be evaluated against the graph metabolite nodes after identifier matching and retention of metabolites measured in at least 30% of samples.

### 2.3 Graph construction and node features

The graph topology is derived from the MetabolicAtlas APIv2 human genome-scale metabolic reconstruction [15], yielding |*V*_*r*_ | = 3,927 reaction nodes and |*V*_*m*_ | = 1,487 metabolite nodes connected by|*ℰ* | = 16,307 directed edges. Edge attributes encode stoichiometric coefficients, with negative values for reactants and positive values for products.

To enable zero-shot generalization, we initialize nodes with domain-specific continuous embeddings rather than one-hot identifiers. Reaction nodes carry feature vectors 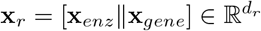, concatenating 768-dimensional EC2Vec [16] functional embeddings (**x**_*enz*_), which encode the catalytic properties of each reaction’s Enzyme Commission number, with normalized gene expression (**x**_*gene*_, TPM). Metabolite nodes use feature vectors 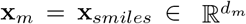 : 384-dimensional ChemBERTa [17] structural embeddings derived from SMILES strings using the DeepChem ChemBERTa-77M-MLM model with mean pooling. Both feature types are projected into a shared latent dimension *d* via learned linear projections:

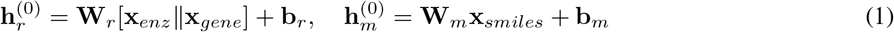

where ∥ denotes concatenation, and 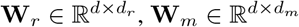 are learnable projection matrices.

We optionally augment node representations with Graph Positional Encodings. For each node, we compute four degree-based features (in-degree, out-degree, total degree, log(degree + 1)) and *K*-step Random Walk Structural Encodings (RWSE) [18]. The resulting (4 + *K*)-dimensional vector (default *K* = 16, yielding 20 features per node) is projected to the hidden dimension and added to the initial node embeddings: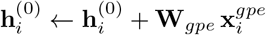.

### 2.4 Cross-modal attention

We employ a bidirectional cross-modal attention mechanism, applied both before the first message-passing layer and after each subsequent GNN layer to re-align the two node-type manifolds. For the metabolite-to-reaction direction, metabolite representations serve as queries while reaction neighbors provide keys and values:

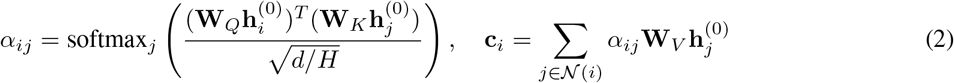

where *H* denotes the number of attention heads. The symmetric reaction-to-metabolite direction is computed analogously with separate projection matrices. The projected context is combined with the input via a residual connection:

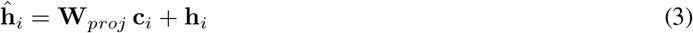

The node embedding is then updated with a second residual connection scaled by *γ* = 0.5:

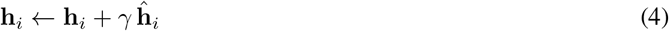

Separate projection matrices are used for each attention layer and each node type.

### 2.5 Relational message passing

We employ a Relational Metabolic Convolution that treats the graph as an explicit bipartite structure with separate learned pathways for each edge direction. At each layer *l*, independent GATv2 [19] attention convolutions process the two edge types in parallel:

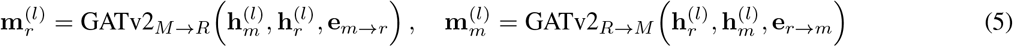

where **e** denotes edge attributes (stoichiometric coefficients) and each GATv2 module uses 4 attention heads. Self-loops are disabled in the neighbor aggregation; instead, each node type maintains a dedicated self-loop processor that computes a self-signal 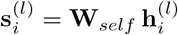. A learned fusion layer combines the neighbor message with the self-signal:

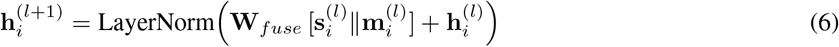

where ∥ denotes concatenation. Messages are passed over *L* layers (default *L* = 3).

When physics-informed training is enabled, a dedicated flux head predicts a scalar flux *v*_*r*_ for each reaction node from the final reaction embeddings:

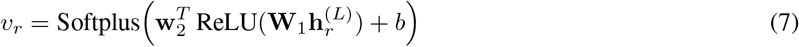

Here **W**_1_, **w**_2_, and *b* are the learnable parameters of the two-layer flux head, and the Softplus activation ensures non-negative flux predictions. These predicted fluxes are used exclusively by the physics-based loss functions (Section 2.7) and do not affect the metabolite concentration predictions in the absence of physics regularization. When flux-gated message passing is additionally enabled, the predicted fluxes modulate information propagation by scaling messages proportionally to predicted reaction activity.

### 2.6 Metabolite-Conditioned Reader

To generalize to unseen metabolites, we replace the standard transductive readout with a query-based attention mechanism we term the Metabolite-Conditioned Reader (MCR). The MCR contains no metabolite-specific parameters; it uses each metabolite’s static structural embedding as a query into the learned reaction representations.

For a target metabolite *m*, the MCR identifies its topological neighbors *N* (*m*) in the metabolic graph and computes a context vector via scaled dot-product attention [20]:

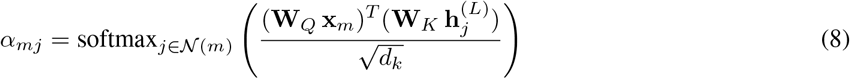

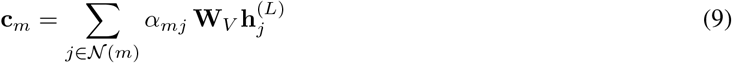

where **W**_*Q*_, **W**_*K*_, and **W**_*V*_ are query, key, and value projections specific to the MCR (distinct from the cross-modal attention projections), *d*_*k*_ is the dimension of the key projection, and **x**_*m*_ is the metabolite’s static structural embedding. The context vector is concatenated with a projected version of the metabolite’s structural embedding and passed through a prediction MLP:

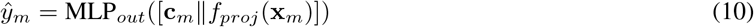

where *f*_*proj*_ is a nonlinear projection (Linear →ReLU) and MLP_*out*_ maps the concatenated representation to a scalar concentration prediction. The MCR further supports multi-hop neighborhood expansion with learnable per-hop attention biases.

### 2.7 Training objectives

The model is trained by minimizing a composite loss function that combines a supervised reconstruction term with a structural alignment auxiliary and three physics-informed regularization terms:

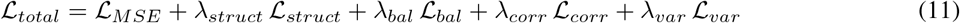

The primary term is a masked mean squared error (MSE) over metabolites with observed measurements in the current sample:

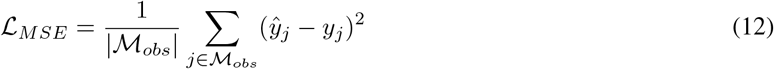

where *ℳ*_*obs*_ denotes the set of metabolites with non-missing concentration values. Missing measurements are common in the CAMP dataset (approximately 180 of 380 mapped metabolites are observed per sample) and are excluded from gradient computation via binary masking.

The *structural alignment* loss is a self-supervised auxiliary that encourages manifold alignment between the GNN’s learned reaction-context representations and the metabolite’s chemical identity. For each metabolite *m*, the messages received from its neighbouring reactions after the final GNN layer are aggregated and passed through a two-layer decoder MLP to predict the metabolite’s SMILES embedding:

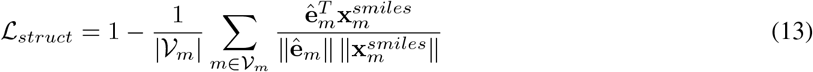

where 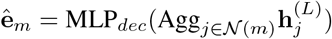 is the decoded embedding. This term requires no additional labels: the frozen ChemBERTa SMILES embeddings serve as targets. It forces the network to encode chemical identity in the reaction neighbourhood representations, which is a prerequisite for the MCR to generalize to unseen metabolites. On a biological graph the functional context of neighbouring enzymes is predictive of molecular identity; on a random graph the context is uninformative and the loss remains high, providing an implicit graph-structure quality signal.

The three physics-informed terms operate on predicted reaction fluxes 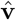 (Section 2.4) and encode biochemical priors without requiring ground-truth flux measurements, drawing on the constraint-based modelling tradition of flux balance analysis [21]. The *mass balance* loss enforces stoichiometric constraints by penalizing net production or consumption at each metabolite node:

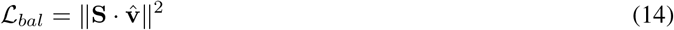

where 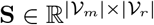 is the stoichiometric matrix derived from edge attributes. At steady state, mass balance requires **S** ·**v** = 0; minimizing *ℒ*_*bal*_ encourages the predicted fluxes to satisfy this constraint. The *gene-flux correlation* loss aligns predicted fluxes with enzyme expression levels, encoding the assumption that highly expressed enzymes tend to catalyse more active reactions:

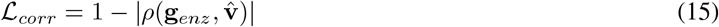

where *ρ* denotes Pearson correlation, | · | denotes absolute value, and **g**_*enz*_ is a per-reaction enzyme activity score derived from the sample’s gene expression by parsing gene-protein-reaction rules compositionally (“and” → min, “or” →max over constituent gene values). The absolute value allows both positive and negative gene–flux correlations to reduce the loss, accommodating reactions where higher enzyme expression may correspond to lower net flux due to reversibility or regulatory effects. Finally, the *flux variance* loss prevents mode collapse by penalizing low spread across the predicted flux distribution:

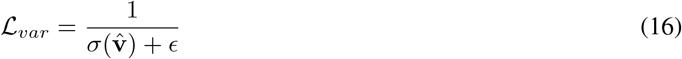

where 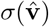 denotes the standard deviation of the predicted flux vector and *ϵ* = 10^*−*6^ is a numerical stability constant. Without this term, the model can trivially minimize *ℒ*_*bal*_ by predicting near-zero fluxes for all reactions. The structural weight *λ*_*struct*_ = 1.0 was held fixed throughout, as the structural alignment term underpins the MCR mechanism rather than acting as a tunable regularization trade-off. The physics weights *λ*_*bal*_, *λ*_*corr*_, and *λ*_*var*_ are treated as hyperparameters and calibrated via grid search (Section 2.7).

### 2.8 Modality ablation study

To determine the contribution of individual biological signals to prediction accuracy, we defined a systematic set of 24 architectural configurations where input channels (metabolite structure **x**_*smiles*_, gene expression **x**_*gene*_, enzyme functional embeddings **x**_*enz*_, and graph positional encodings **x**_*gpe*_) were independently ablated by replacement with Gaussian noise *N* (0, **I**) of matching dimensionality. Topological dependence was similarly assessed by substituting the biological adjacency matrix **A** with a density-matched Erdős–Rényi random graph **A**_*rand*_ [22], isolating the contribution of metabolic pathway structure from the general capacity of graph propagation. The search space further encompassed the presence or absence of cross-modal attention, the MCR, and GPE (including a randomized-GPE control to separate learned structural signal from added parameter capacity). We additionally tested a mean-subtraction preprocessing variant where per-sample gene expression is centered prior to input. Each configuration was evaluated using 5-fold cross-validation. An XGBoost [23] baseline consisting of one independent gradient boosting model per metabolite was trained under the same data splits as a non-graph reference. Separately, the three physics loss weights were calibrated via a grid search over 60 combinations (*λ*_*bal*_ ∈ {0.0, 0.1, 0.5, 1.0, 2.0}, *λ*_*corr*_ ∈ {0.0, 0.1, 0.5, 1.0}, *λ*_*var*_ ∈ {0.0, 1.0, 2.0}) using the full model architecture with a fixed random seed; optimal weights were carried forward to the LOMO evaluation.

### 2.9 Leave-One-Metabolite-Out validation

Generalization capacity to truly unseen metabolites is evaluated under a Leave-One-Metabolite-Out (LOMO) cross-validation protocol. For a designated target metabolite *m*, all ground-truth concentration values *y*_*·*,*m*_ are masked during training; the model must predict these values using only the metabolite’s structural embedding and its position in the graph, without ever having observed a single training example for that compound. Eligible metabolites are those with at least 20 observed samples in the dataset. From this set, a balanced subset of 50 metabolites is selected by stratifying across neighbourhood coverage quantiles (5 quantile bins), ensuring evaluation includes both highly connected metabolic hubs and sparsely connected compounds. For each LOMO fold, a fresh model is trained from random initialization for 300 epochs (learning rate = 1 × 10^*−*3^, batch size 4 with gradient accumulation over 4 steps) using a 70/30 train/test split of cell lines.

### 2.10 Implementation details

All models were implemented in PyTorch Geometric [24] and trained using the Adam optimizer [25] (*η* = 7.5 × 10^*−*4^, weight decay = 5 × 10^*−*4^). The learning rate was reduced on plateau (factor 0.5, patience 5 epochs). We used a batch size of 1, processing the full heterogeneous graph as a single sample per forward pass. Training persisted for a maximum of 300 epochs with early stopping (patience = 10 epochs, monitoring validation loss). The default hidden dimension was *d* = 128 with *L* = 3 GNN layers and dropout *p* = 0.2.

R^2^ and RMSE are computed over the pooled set of all observed (sample, metabolite) pairs. This pooled metric includes both between-metabolite and within-metabolite variance; we report per-metabolite R^2^ distributions where available (Section 3.2). The XGBoost baseline trains one independent regressor per metabolite using the same gene-expression features, with predictions pooled identically.

### 2.11 Comparison with existing methods

To contextualize GAZE’s zero-shot performance, we asked how well existing methods can predict metabolite levels from gene expression on the CAMP dataset. The common evaluation is: for each metabolite, compute a predicted score across 867 cell lines using only gene expression (and, for UnitedMet, other metabolites’ measurements), then measure the Spearman rank correlation between predicted and measured concentrations. Because each method produces a different output type (activity scores, task scores, or predicted ranks), Spearman *ρ* provides a scale-invariant metric that enables direct comparison.

We evaluated three methods spanning two paradigms. Knowledge-based methods (scCellFie, MEBOCOST) use curated enzyme–metabolite databases to compute metabolite activity scores from gene expression, without any training. Supervised imputation (UnitedMet) learns the joint gene–metabolite distribution from paired measurements and imputes held-out metabolites. In contrast, GAZE operates in a zero-shot setting: the target metabolite is entirely excluded from training, and prediction relies on graph topology and chemical representations.

scCellFie [7] scores metabolic tasks by evaluating gene-protein-reaction (GPR) rules against expression data. Each task corresponds to a metabolic function (e.g., “Glutamine degradation”) and is scored as the minimum across its constituent reactions, where each reaction score is the mean expression of its associated genes. We loaded the human scCellFie database (218 tasks, 787 reactions, 931 genes) and scored all tasks across the 867 CAMP cell lines. Because scCellFie expects raw counts and normalizes internally to counts per 10,000 (CP10k), we rescaled the TPM expression matrix to CP10k and disabled the internal normalization step. We mapped task names to CAMP metabolites by extracting the metabolite portion of each task name and matching against the 1,152 CAMP metabolite names via case-insensitive string matching. This yielded 142 task-metabolite pairs with at least 20 valid samples.

MEBOCOST [8] estimates metabolite activity from enzyme expression using a curated database mapping metabolites to producing and consuming enzymes via HMDB identifiers. For each metabolite, activity is: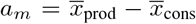, where 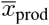 and 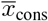 are the mean expression of producing and consuming enzyme genes, respectively. We matched 260 MEBOCOST metabolites to the CAMP panel by normalizing HMDB accession formats.

UnitedMet [5] is a Bayesian matrix factorization method that models the joint gene–metabolite distribution using a Plackett-Luce ranking likelihood over a low-rank latent space (**Z** = **WH**, *K* = 30), with inference via stochastic variational inference (4,000 steps). It predicts metabolite ranks rather than absolute concentrations. UnitedMet was designed for *sample-holdout* imputation, in which all metabolite features are observed during training and the model imputes values for new patient samples. Our evaluation protocol tests a different capability: *metabolite-holdout* prediction, in which entire metabolite features are masked during training. This is not the task UnitedMet was designed for, and its performance under this protocol should be interpreted accordingly.

We applied UnitedMet to the CAMP dataset using the 5,000 most variable genes and 262 metabolites with ≥ 50% sample coverage. To evaluate metabolite-holdout performance analogous to GAZE’s LOMO protocol, we performed 10-fold cross-validation over metabolite features: in each fold, one tenth of the metabolites (~ 26) were masked and the model was trained on all remaining metabolites plus all genes, then posterior rank predictions were generated (500 draws). Notably, even under this metabolite-holdout protocol, UnitedMet retains access to the measured concentrations of all non-masked metabolites during training. This information is unavailable to GAZE, scCellFie, or MEBOCOST.

### 2.12 External validation on independent tissue cohort

To test whether zero-shot predictions generalize beyond the training domain, we deployed the trained GAZE model on the UnitedMet ccRCC cohort without any fine-tuning or domain adaptation. The model was trained on all 867 CAMP cell lines using the best configuration from the LOMO evaluation (relational convolution, cross-modal attention, MCR, physics constraints). At inference, external gene expression profiles were passed through the same metabolic graph and processing pipeline used during training.

For each of the 214 metabolites retained for evaluation, we computed the Spearman rank correlation between GAZE predictions and measured concentrations across all tissue samples in which that metabolite was detected. As baselines, we applied MEBOCOST and scCellFie to the same external RNA-seq data using their respective enzyme–metabolite databases, yielding 188 and 123 metabolites, respectively. Because UnitedMet is a supervised method that requires paired gene–metabolite measurements for training, it cannot be applied in a single forward pass like GAZE. Instead, we ran UnitedMet on the same RC18+RC20 data using 10-fold metabolite-holdout cross-validation (5,000 most variable genes, *K* = 30, 4,000 SVI steps per fold): in each fold, one tenth of the metabolites were masked during training, and the model predicted their ranks from the remaining metabolites and all genes. This yielded 532 metabolites. As in the CAMP comparison, this applies UnitedMet to a metabolite-holdout task rather than its intended sample-holdout task. The configuration is otherwise maximally favourable: UnitedMet is trained and evaluated on the same ccRCC tissue cohort for which it was originally developed, and retains access to all non-held-out metabolite measurements during training.

## 3 Results

### 3.1 GAZE: a graph attention architecture for zero-shot metabolite estimation

Figure 1 summarises the GAZE architecture. The model operates on a bipartite metabolic network in which reaction nodes carry concatenated EC2Vec enzyme embeddings and sample-specific gene expression (TPM), while metabolite nodes are initialised from ChemBERTa SMILES embeddings. Three layers of Relational Metabolic Convolution propagate information along reaction-to-metabolite and metabolite-to-reaction edges via independent GATv2 attention heads, interleaved with bidirectional cross-modal attention that re-aligns the two node-type manifolds after each round of message passing. Because the readout must generalise to metabolites absent from training, the standard per-node prediction head is replaced by the Metabolite-Conditioned Reader (MCR): a query-based attention mechanism that uses each metabolite’s SMILES embedding as a query into the contextualised reaction representations of its graph neighbours, producing a scalar concentration prediction through shared weights with no metabolite-specific parameters. An auxiliary flux head predicts non-negative reaction fluxes whose product with the stoichiometric matrix provides a physics-informed mass-balance regulariser during training.

**Figure 1:**
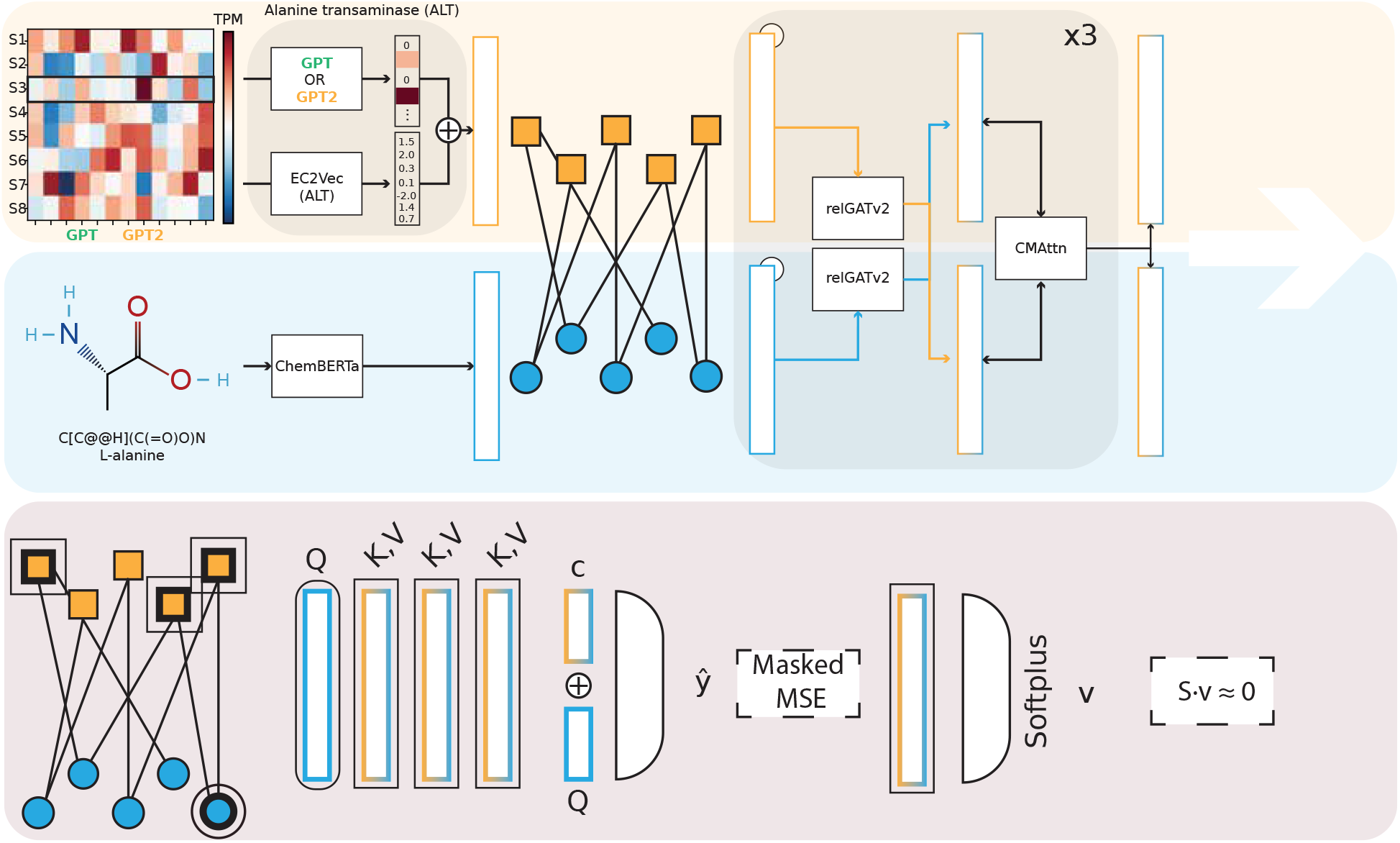
Overview of the GAZE architecture. Reaction nodes (orange squares) are initialised with concatenated EC2Vec enzyme embeddings and sample-specific gene expression (TPM); metabolite nodes (blue circles) are initialised from ChemBERTa SMILES embeddings. Both modalities are projected into a shared latent space and connected through a bipartite metabolic graph. Three layers of relational GATv2 convolution propagate information along separate reaction-to-metabolite and metabolite-to-reaction edges, interleaved with bidirectional cross-modal attention (CMAttn) that re-aligns the two node-type representations. At the readout stage (bottom), the Metabolite-Conditioned Reader (MCR) uses each metabolite’s SMILES embedding as a query (*Q*) into the contextualised reaction embeddings of its graph neighbours (keys *K*, values *V*), producing a scalar concentration prediction *ŷ* through shared weights with no metabolite-specific parameters. An auxiliary flux head applies a Softplus activation to predict non-negative reaction fluxes **v**, regularised by the stoichiometric mass-balance constraint *S***v** ≈ 0.

### 3.2 Modality ablation identifies graph topology and gene expression as the dominant predictive signals

To determine which input signals and structural priors drive metabolite prediction, we evaluated a set of architectural configurations across 5-fold cross-validation (**Table 1**). The search space encompassed input modalities (gene expression, EC2Vec enzyme embeddings, ChemBERTa SMILES embeddings, graph positional encodings) and graph topology (biological vs. randomized). As a non-graph reference, we trained an XGBoost baseline consisting of one independent model per metabolite under the same data splits, which achieved R^2^ = 0.835 ± 0.003. All pairwise comparisons report Welch’s *t*-tests across the five folds.

**Table 1:** Modality ablation results. All configurations use the biological metabolic graph. R^2^ and RMSE are averaged over 5 folds (± std). GAZE denotes the full architecture (cross-modal attention, residual connections, MCR, 50% SMILES masking).

| Configuration | $R^2$ (↑) | RMSE (↓) |
| --- | --- | --- |
| XGBoost (per-metabolite) | $0.835 \pm 0.003$ | $1.668 \pm 0.025$ |
| GAZE (mean-subtracted) | $0.832 \pm 0.012$ | $1.798 \pm 0.068$ |
| All inputs (transductive GATv2) | $0.824 \pm 0.011$ | $1.827 \pm 0.059$ |
| Gene expression only | $0.823 \pm 0.008$ | $1.850 \pm 0.040$ |
| EC2Vec + gene expression | $0.821 \pm 0.011$ | $1.846 \pm 0.070$ |
| GAZE | $0.816 \pm 0.013$ | $1.884 \pm 0.068$ |
| GAZE + GPE | $0.814 \pm 0.009$ | $1.892 \pm 0.044$ |
| Graph Only (bio topology) | $0.711 \pm 0.015$ | $2.359 \pm 0.068$ |
| SMILES only | $0.401 \pm 0.004$ | $3.399 \pm 0.032$ |
| Enzyme embeddings only | $0.385 \pm 0.004$ | $3.420 \pm 0.033$ |
| GPE only | $0.377 \pm 0.005$ | $3.466 \pm 0.032$ |

The single largest factor in the ablation was biological graph topology. A GNN operating on the metabolic graph with noise-initialized node features achieved R^2^ = 0.711± 0.015, a 39.8% improvement over the same architecture on a density-matched random graph (R^2^ = 0.509± 0.015; *p <* 0.0001; **Supplementary Figure 6**). This gap was most pronounced for feature-free configurations; when strong node features were present, random and biological graphs converged (e.g., GAZE: 0.816 vs. 0.794, Δ = 0.022), suggesting that rich features can partially compensate for missing topological structure. Adding gene expression features to the biological graph further increased performance to R^2^ = 0.823 ± 0.008 (+15.6%). Gene expression alone on a random graph still reached R^2^ = 0.800, indicating that gene–metabolite statistical associations carry substantial predictive signal independently of network structure.

GAZE achieved R^2^ = 0.816 ± 0.013, a similar performance compared to a transductive GATv2 encoder with the same input modalities but without the generalization components (R^2^ = 0.824 ± 0.011; *p* = 0.35). Notably, the transductive encoder cannot predict metabolites absent from training. GAZE’s cross-modal attention, MCR, and 50% SMILES masking introduce zero-shot capacity at no significant cost to in-distribution accuracy.

No additional modality beyond graph topology and gene expression provided a statistically significant improvement on the standard prediction task. Concatenating EC2Vec enzyme embeddings with gene expression (R^2^ = 0.821 ± 0.011) did not improve over gene expression alone (R^2^ = 0.823; *p* = 0.80), and incorporating the full zero-shot architecture (SMILES embeddings, cross-modal attention, and MCR) caused no significant decrease relative to the enzyme-gene configuration (*p* = 0.58). In isolation, both EC2Vec (R^2^ = 0.385) and SMILES (R^2^ = 0.401) performed comparably to noise, confirming that these embeddings carry limited predictive information when not grounded by gene expression. We similarly tested Graph Positional Encodings (GPE), which provide explicit degree and random-walk structural features. GPE showed no additive benefit when added to GAZE (R^2^ = 0.814± 0.009 vs. 0.816 ± 0.013; *p* = 0.87). A randomized-GPE control that preserves the additional parameters while destroying structural content performed equivalently (R^2^ = 0.811 ± 0.014; *p* = 0.73 vs. real GPE; **Supplementary Figure 7**), ruling out both redundant structural signal and insufficient parameter capacity as explanations. GPE in isolation (R^2^ = 0.377) confirmed that degree and random-walk statistics alone carry negligible metabolic information.

Although EC2Vec and SMILES embeddings do not improve standard prediction, they serve a different function: they are the mechanism through which the model can generalize to metabolites absent from the training set. The MCR uses each metabolite’s SMILES embedding as a query into the learned reaction representations, and EC2Vec provides transferable functional descriptors that are not tied to specific metabolite identities. Without these modalities, the architecture reduces to a transductive model that can only predict metabolites seen during training, equivalent in scope to the XGBoost baseline. A mean-subtracted preprocessing variant of GAZE achieved R^2^ = 0.832 ± 0.012, approaching the XGBoost baseline; however, as the physics calibration and LOMO experiments were conducted under standard preprocessing, we retained GAZE with standard normalization (R^2^ = 0.816) as the reference architecture throughout (**Supplementary Figure 8**).

Next, we calibrated the physics loss weights via a grid search over 60 combinations on a fixed evaluation seed. The best configuration (*λ*_*bal*_ = 0.5, *λ*_*corr*_ = 0.1, *λ*_*var*_ = 1.0) achieved test R^2^ = 0.809 versus 0.803 for the no-physics baseline, with the top-10 configurations clustered within R^2^ = 0.804–0.809. These weights were carried forward to the LOMO evaluation.

### 3.3 Zero-shot generalization via Leave-One-Metabolite-Out evaluation

To assess whether the architectural components identified above enable genuine zero-shot prediction, we evaluated 10 configurations under the LOMO protocol (Section 2.6). Each configuration was trained from scratch for each of 50 held-out metabolites, selected by stratified sampling across neighbourhood coverage quantiles (**Supplementary Figure 9**). **Table 2** summarizes the results, organized into a cumulative build-up, ablations that remove key components from GAZE, and additional variants.

**Table 2:** Leave-One-Metabolite-Out (LOMO) zero-shot evaluation across 10 architectural configurations. Each configuration was trained from scratch on 50 held-out metabolites (300 epochs per fold, 70/30 train/test split). Configurations are grouped by component family. “% R^2^ *>* 0” indicates the fraction of held-out metabolites for which the model outperforms a constant mean predictor. MCR = Metabolite-Conditioned Readout; GPE = Graph Positional Encodings.

| Configuration | Median $R^2$ | % $R^2 > 0$ | # $R^2 > 0.5$ | Max $R^2$ |
| --- | --- | --- | --- | --- |
| *Cumulative build-up* |  |  |  |  |
| Homogeneous GATv2 | -0.72 | 16 | 1 | 0.55 |
| + Bipartite message passing + MCR | -0.53 | 18 | 1 | 0.74 |
| + Residual connections | -0.52 | 22 | 1 | 0.61 |
| GAZE (+ cross-modal attn + SMILES mask.) | -0.35 | 24 | 4 | 0.76 |
| *Ablations (components removed from GAZE)* |  |  |  |  |
| Cross-modal attention only (no MCR) | -0.82 | 20 | 2 | **0.86** |
| Bipartite GNN + GPE (no MCR) | -0.63 | 14 | 1 | 0.83 |
| *GAZE variants* |  |  |  |  |
| GAZE + multi-hop readout | -0.41 | 22 | 4 | 0.79 |
| GAZE + multi-hop readout + GPE | -0.41 | 22 | 2 | 0.76 |
| *Physics-informed* |  |  |  |  |
| GAZE + physics | **-0.34** | **30** | **5** | 0.78 |
| GAZE + physics + multi-hop + GPE | -0.60 | 18 | 1 | 0.81 |

All configurations produced negative median R^2^, reflecting the inherent difficulty of predicting a metabolite the model has never observed during training. An R^2^ of zero corresponds to predicting each metabolite’s mean concentration, so negative values indicate predictions worse than this constant baseline. However, we observed a consistent improvement across the cumulative build-up (**Figure 2**). The homogeneous GATv2 baseline, which uses a transductive readout forced into zero-shot mode, achieved median R^2^ = −0.72 with only 16% of metabolites producing positive R^2^. Replacing the homogeneous message passing with a bipartite Relational HeteroConv and MCR readout improved the median to − 0.53 (+0.19), and adding residual connections further narrowed the deficit to − 0.52 while increasing the positive-R^2^ fraction to 22%. The complete GAZE architecture, combining relational convolution, residual connections, cross-modal attention, and the MCR, achieved median R^2^ = −0.35 with 24% positive R^2^ and four metabolites exceeding R^2^ *>* 0.5.

**Figure 2:**
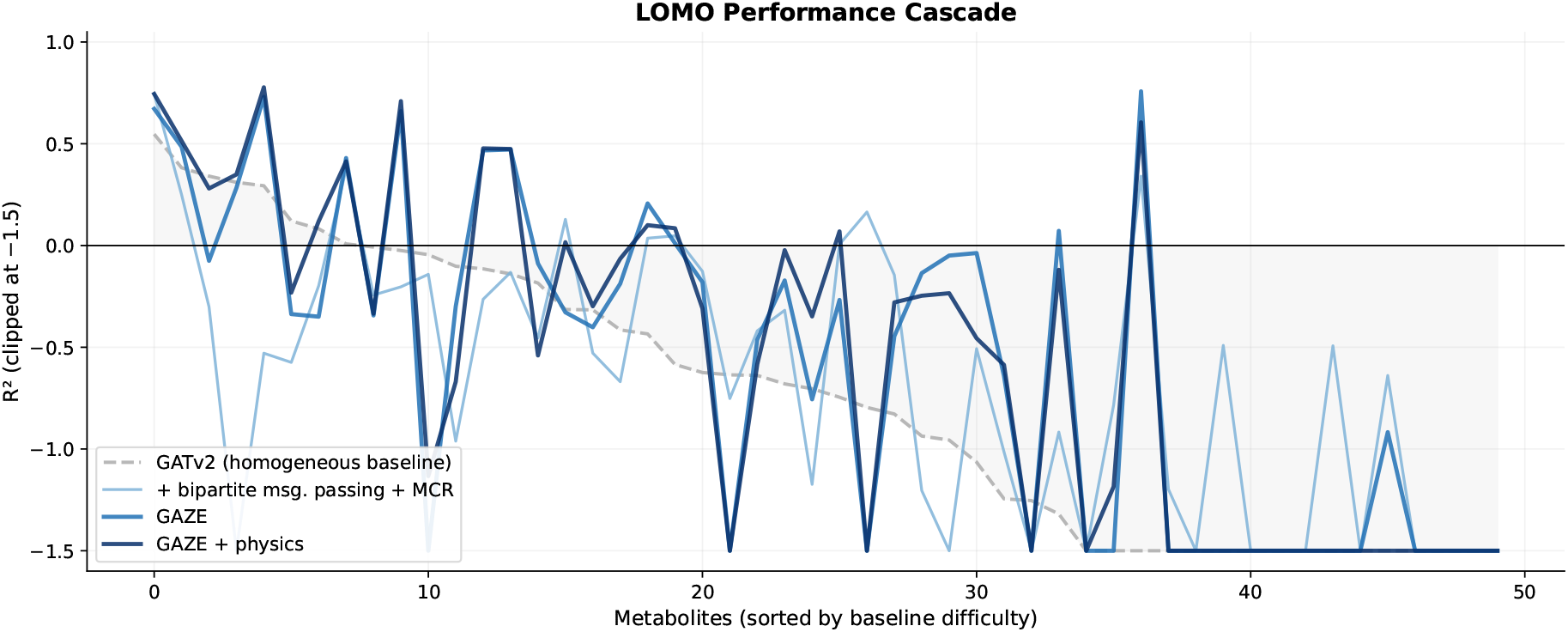
Per-metabolite LOMO R^2^ across four architectural stages, sorted by baseline difficulty (descending). Each line traces one of the 50 held-out metabolites across configurations: GATv2 baseline (grey), + HeteroConv (light blue), GAZE (medium blue), and GAZE + physics (dark blue).

Adding physics-informed training (flux gating and mass balance projection) to GAZE produced the best overall configuration: median R^2^ = −0.34, 30% positive R^2^, and five metabolites above R^2^ *>* 0.5. While the median R^2^ improvement over GAZE was modest (Δ = +0.01), physics regularization increased the fraction of positive-R^2^ metabolites by 6 percentage points, suggesting that biochemical constraints broaden the set of metabolites for which the model learns transferable representations rather than dramatically improving any single prediction.

The per-metabolite R^2^ distributions exhibited heavy tails: the best individual metabolite in the physics-informed model reached R^2^ = 0.78, and multiple architectures produced individual R^2^ values exceeding 0.75. In contrast, a small number of metabolites produced R^2^≪ −10, dominating the mean but not the median. This high inter-metabolite variance motivated the mechanism analysis below, which investigates which metabolite-level properties govern zero-shot predictability.

Ablation experiments confirmed that both the MCR and cross-modal attention are jointly necessary. Removing the MCR and retaining only cross-modal attention degraded median R^2^ to −0.82 (the worst among all configurations) despite producing the highest single-metabolite R^2^ (0.86). This indicates that cross-modal attention produces high-magnitude but poorly calibrated representations without the MCR’s query-based readout, and that the two components must work together for stable zero-shot performance. Similarly, extending GAZE with additional physics terms (GAZE + physics extended) degraded performance relative to the standard physics variant (median −0.60 vs. −0.34), suggesting that excessive regularization can reduce generalization performance.

To understand why certain metabolites are predictable while others fail catastrophically, we correlated per-metabolite LOMO R^2^ with five candidate features: chemical support (maximum Tanimoto similarity to measured metabolites), enzyme signal strength (mean expression variance of neighbouring enzymes), graph degree, molecular size (carbon count), and target variability (interquartile range of measured concentrations across 867 cell lines). Of the 50 LOMO metabolites, 15 lacked a resolved SMILES string; these consistently produced the most extreme failures, with the majority achieving R^2^ *<*− 2 (**Figure 3A**). Among the 35 metabolites with resolved SMILES, target variability emerged as the dominant predictor of zero-shot success: the interquartile range of measured concentrations correlated strongly with R^2^ (Spearman *ρ* = 0.66, *p <* 10^*−*4^; **Figure 3B**). Molecular size showed a secondary negative effect (carbon count: *ρ* = −0.37, *p* = 0.03; **Figure 3C**), while chemical similarity (*ρ* = 0.17, *p* = −0.34), graph degree (*ρ* = 0.18, *p* = 0.30), and enzyme signal (*ρ* = 0.07, *p* = 0.72) exhibited no significant relationship. Notably, chemical similarity to the training metabolome does not determine success among resolved metabolites. Rather, the model’s ability to generalize depends on whether the target metabolite carries a learnable signal in the expression data.

**Figure 3:**
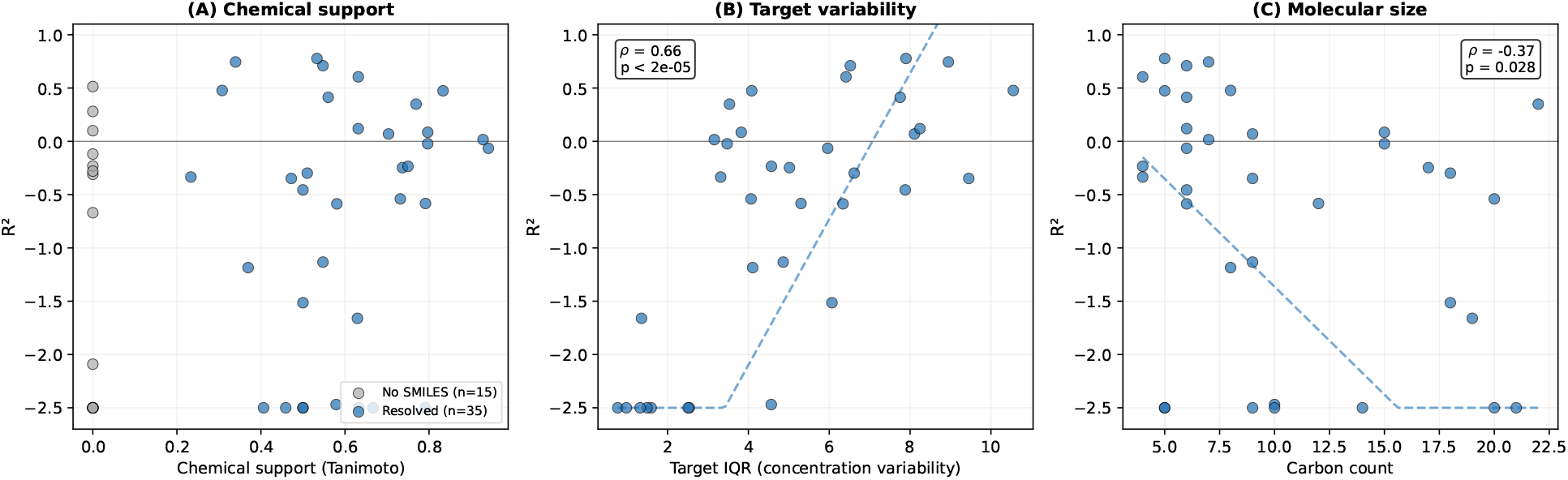
Determinants of zero-shot predictability. **(A)** Chemical support (Tanimoto similarity) vs LOMO R^2^; metabolites without resolved SMILES shown in grey. **(B)** Target variability (IQR of measured concentrations) vs R^2^ for resolved metabolites. **(C)** Molecular size (carbon count) vs R^2^ for resolved metabolites. Dashed lines show linear regression trends; R^2^ values clipped at −2.5.

Eight metabolites exceeded R^2^ *>* 0.4 in both the GAZE and GAZE + physics configurations: hypoxanthine (R^2^ = 0.73/0.77), 1-methylnicotinamide (0.67/0.74), L-histidine (0.71/0.71), L-threonine (0.61/0.76), sedoheptulose 7-phosphate (0.51/0.51), ribitol (0.47/0.47), acetaminophen (0.48/0.48), and L-cystine (0.41/0.43). In addition to the statistical prerequisites identified above, these metabolites share two structural properties. They occupy an intermediate graph degree of 2–8, sufficiently connected for neighboring nodes to constrain the prediction but not so central that their removal disrupts the local subgraph. They also sit at fixed stoichiometric positions within linear or near-linear pathway segments: 1-methylnicotinamide is the terminal product of nicotinamide N-methyltransferase (NNMT) acting on nicotinamide [26], L-cystine is the direct oxidation product of cysteine via a single reversible reaction, and sedoheptulose 7-phosphate occupies a fixed position between ribulose 5-phosphate and fructose 6-phosphate in the non-oxidative pentose phosphate pathway [27]. In each case, the flanking metabolites that remain in training tightly constrain the held-out node through stoichiometric relationships, suggesting that the model generalizes primarily through local pathway interpolation rather than globally transferable gene-to-metabolite rules. In contrast, we identified two catastrophic failure modes. The most severe failures (R^2^ ≪ −10) occurred among the 15 metabolites (30% of the LOMO set) lacking a resolved SMILES string, including creatine phosphate (R^2^ = −22.8), 2,3-bisphosphoglycerate (−22.4), phenylacetylglycine (−21.6), and betaine aldehyde (−19.4). Without a chemical embedding the MCR cannot query the graph (median R^2^ = − 0.67 vs. 0.34 for resolved metabolites). A second mode affected large, structurally complex molecules even when SMILES were available: cortisol (R^2^ = −15.0, 21 carbons), deoxyadenosine (−9.8, 10 carbons), and prostaglandin E2 (−6.4, 20 carbons), consistent with the negative correlation between molecular size and R^2^ reported above. Hub metabolites with graph degree *>* 10 behaved differently. Rather than failing catastrophically, they clustered at near-zero R^2^ (−0.54 to +0.08) regardless of chemical support, neither attaining the positive R^2^ of well-placed intermediate-degree metabolites nor incurring the large errors of the two failure modes above.

To determine whether the physics-informed GAZE variant (the best-performing configuration) learns meaningful representations even for metabolites with negative R^2^, we examined the raw per-sample predictions and targets for the subset of LOMO metabolites with available sample-level predictions. Among metabolites with negative R^2^, 77% exhibited positive Pearson correlation with the true concentrations, and 62% reached statistical significance (*p <* 0.05). Only 3 of 20 tested metabolites showed genuinely negative or zero correlation. The negative R^2^ arises primarily from a calibration deficit: because the model has never observed the target metabolite’s concentration distribution during training, its predictions are shifted in mean (median |bias| = 1.3 units) and compressed in dynamic range (median scale ratio = 0.55). Post-hoc z-score recalibration (rescaling predictions to match the target mean and variance) improved median R^2^ from −0.38 to −0.30 and converted individual metabolites from negative to positive R^2^ (e.g., one metabolite improved from R^2^ = −0.05 to +0.17 with Pearson *r* = 0.59). This pattern was consistent across all architectures tested. These results indicate that the model captures biologically meaningful covariation for the majority of held-out metabolites, but cannot anchor predictions to the correct absolute concentration scale because no target-metabolite measurements are available during training.

### 3.4 Comparison with existing methods

We compared GAZE’s zero-shot predictions against two knowledge-based methods and one supervised imputation method (Section 2.9; **Figure 4A**). Because each method covers a different subset of metabolites (MEBOCOST: 260, UnitedMet: 262, scCellFie: 142, GAZE LOMO: 50), we report both per-method distributions and pairwise comparisons on overlapping metabolites. GAZE achieved a median Spearman *ρ* = 0.417 across 50 LOMO metabolites, with 82% showing positive correlation. MEBOCOST achieved a median Spearman *ρ* = 0.040 across 260 metabolites, with only 53% showing positive correlation. scCellFie achieved median *ρ* = 0.158 across 142 task-metabolite pairs with 61% positive correlations. On the 8 metabolites where both GAZE and scCellFie could be evaluated, GAZE produced the stronger absolute correlation for all 8 (**Figure 4B**). On the 32 metabolites where both MEBOCOST and GAZE LOMO predictions were available, GAZE produced the stronger correlation for 24 of 32 (**Figure 4C**).

**Figure 4:**
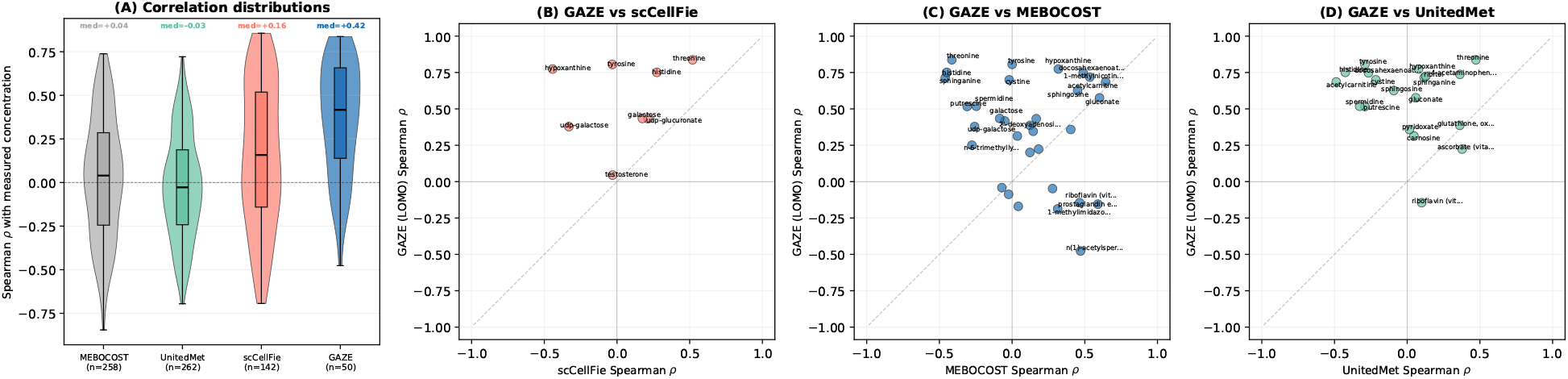
Comparison of GAZE against existing methods on CAMP data. **(A)** Distribution of per-metabolite Spearman *ρ* for MEBOCOST (*n* = 260), UnitedMet (*n* = 262), scCellFie (*n* = 142), and GAZE LOMO (*n* = 50). Dashed line marks *ρ* = 0. **(B)** GAZE vs scCellFie on 8 overlapping metabolites. **(C)** GAZE vs MEBOCOST on 32 overlapping LOMO metabolites. **(D)** GAZE vs UnitedMet on 19 overlapping metabolites.

UnitedMet, a supervised Bayesian imputation method that has access to all other metabolites’ measurements during training, achieved a median *ρ* = −0.028 across 262 metabolites in 10-fold metabolite-holdout cross-validation, with only 47% positive correlations (**Figure 4A**). On the 19 metabolites where both GAZE and UnitedMet could be evaluated, GAZE produced the stronger correlation for 18 (**Figure 4D**). We stress that this metabolite-holdout evaluation tests a capability outside UnitedMet’s design scope: the method was built for sample-holdout imputation, where all metabolite identities are known at training time. Its poor performance here does not indicate a flaw in the method, but rather demonstrates that the statistical covariance structure captured by matrix factorization does not, on its own, enable generalization to metabolites absent from training. This is a zero-shot prediction problem [6]: without side information describing the target metabolite, a model that relies solely on co-occurrence statistics cannot generalize to unseen items. The structural priors used by GAZE (metabolic network topology and chemical representations) provide information that latent factorization cannot recover from gene–metabolite covariance alone.

### 3.5 External validation on independent tissue cohort

To test whether the learned representations transfer across biological domains, we deployed the trained GAZE model on the RC18 and RC20 sub-cohorts of the UnitedMet ccRCC cohort (220 tumour tissue samples; Section 2.12) without any fine-tuning (**Figure 5**).

**Figure 5:**
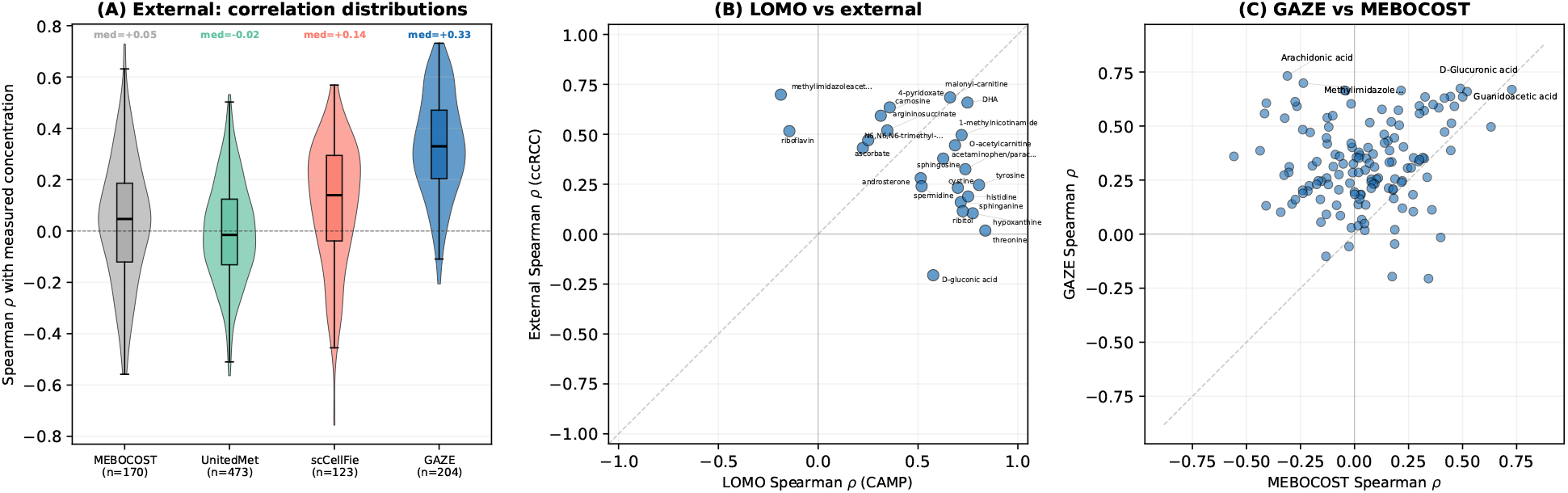
External validation on the UnitedMet ccRCC tissue cohort (RC18 + RC20, *n* = 220 samples). GAZE was trained on CAMP cell lines only; no fine-tuning was applied. **(A)** Distribution of per-metabolite Spearman *ρ* for MEBOCOST (*n* = 172), UnitedMet (*n* = 473), scCellFie (*n* = 123), and GAZE (*n* = 204). **(B)** Concordance between per-metabolite Spearman *ρ* on CAMP (LOMO) and external ccRCC for overlapping metabolites. **(C)** GAZE vs MEBOCOST pairwise on overlapping external metabolites.

**Figure 6:**
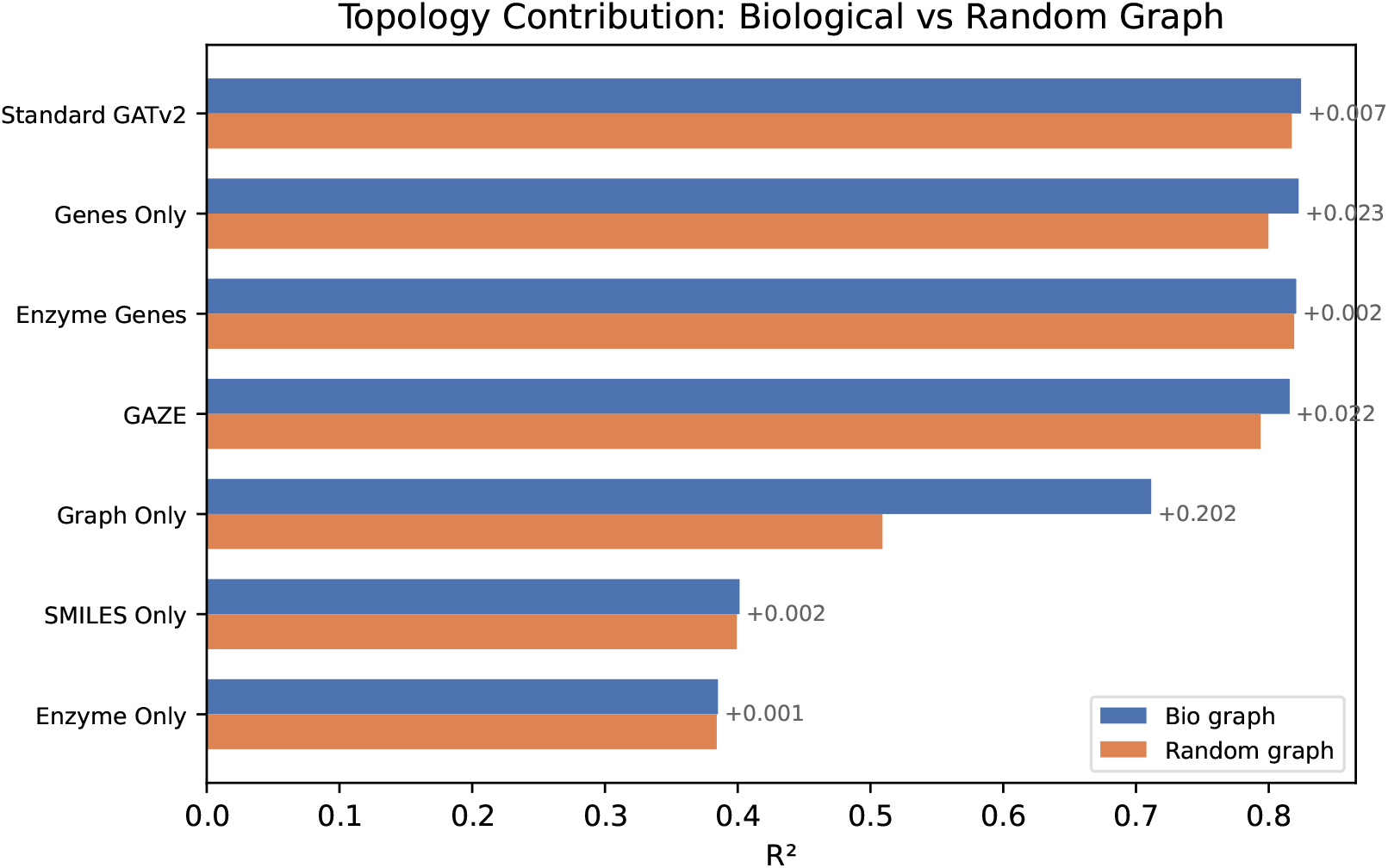
Topology contribution across modality configurations. Paired horizontal bars show R^2^ on the biological metabolic graph (blue) versus a density-matched random graph (orange) for each input modality.

**Figure 7:**
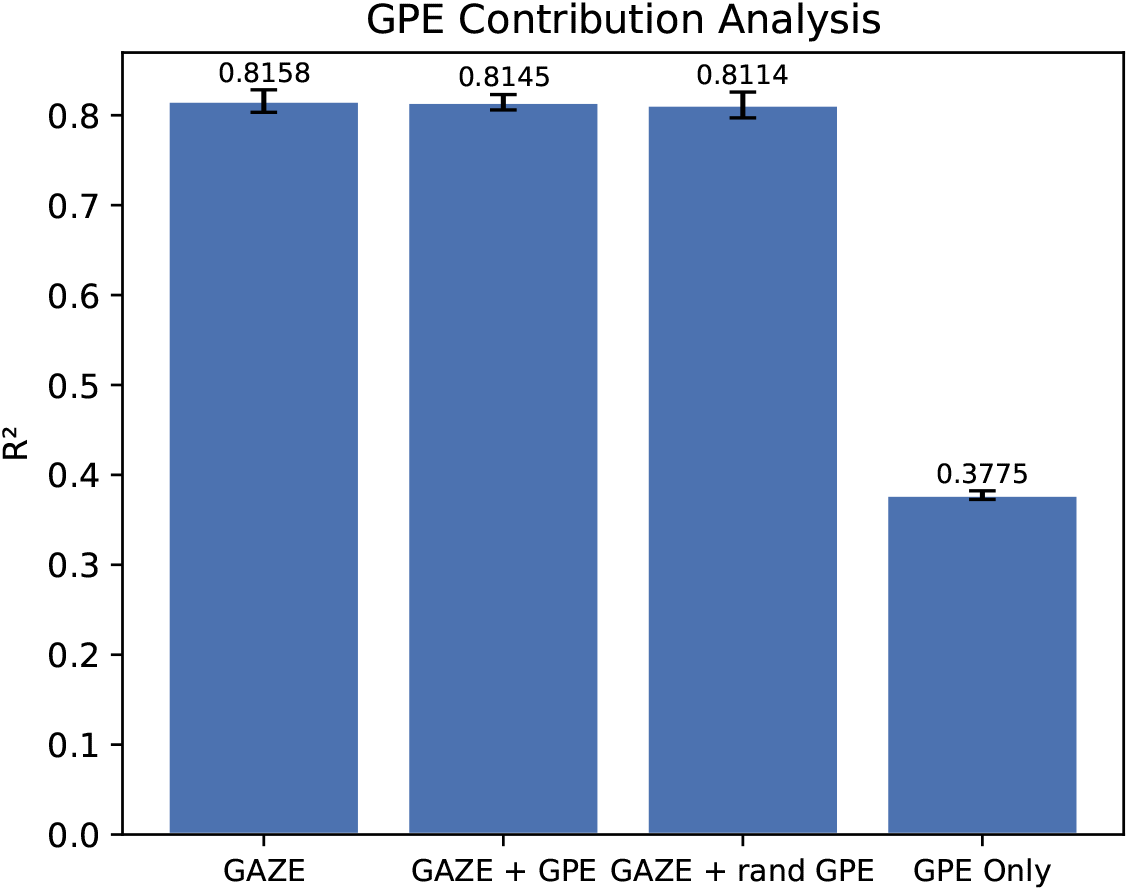
Graph Positional Encoding (GPE) contribution analysis. GAZE compared against three conditions: real GPE, randomized GPE (parameter-count control), and GPE as sole input.

**Figure 8:**
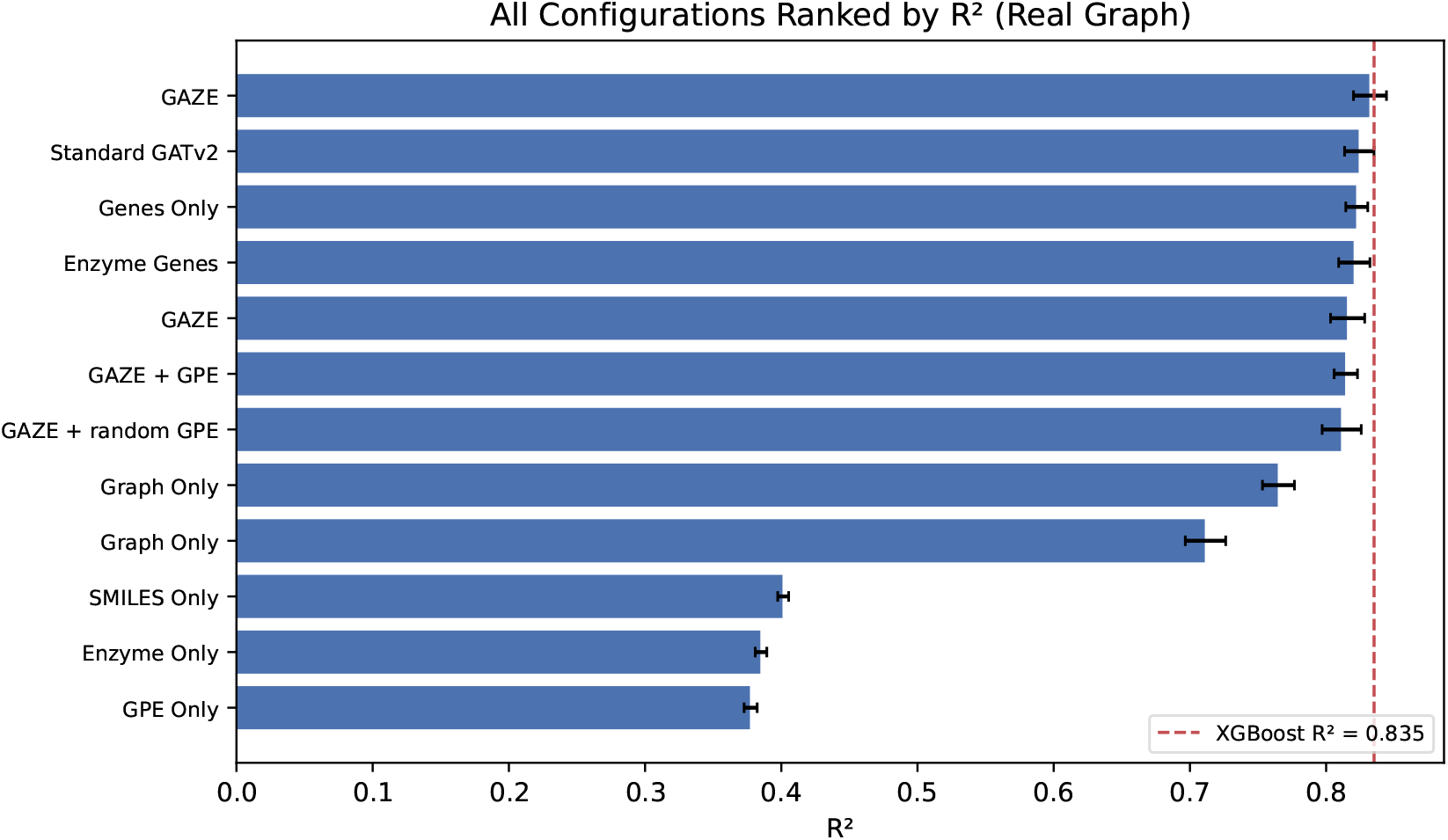
All real-graph configurations ranked by mean R^2^ across 5 folds. Error bars show standard deviation. The dashed red line marks the per-metabolite XGBoost baseline (R^2^ = 0.835).

**Figure 9:**
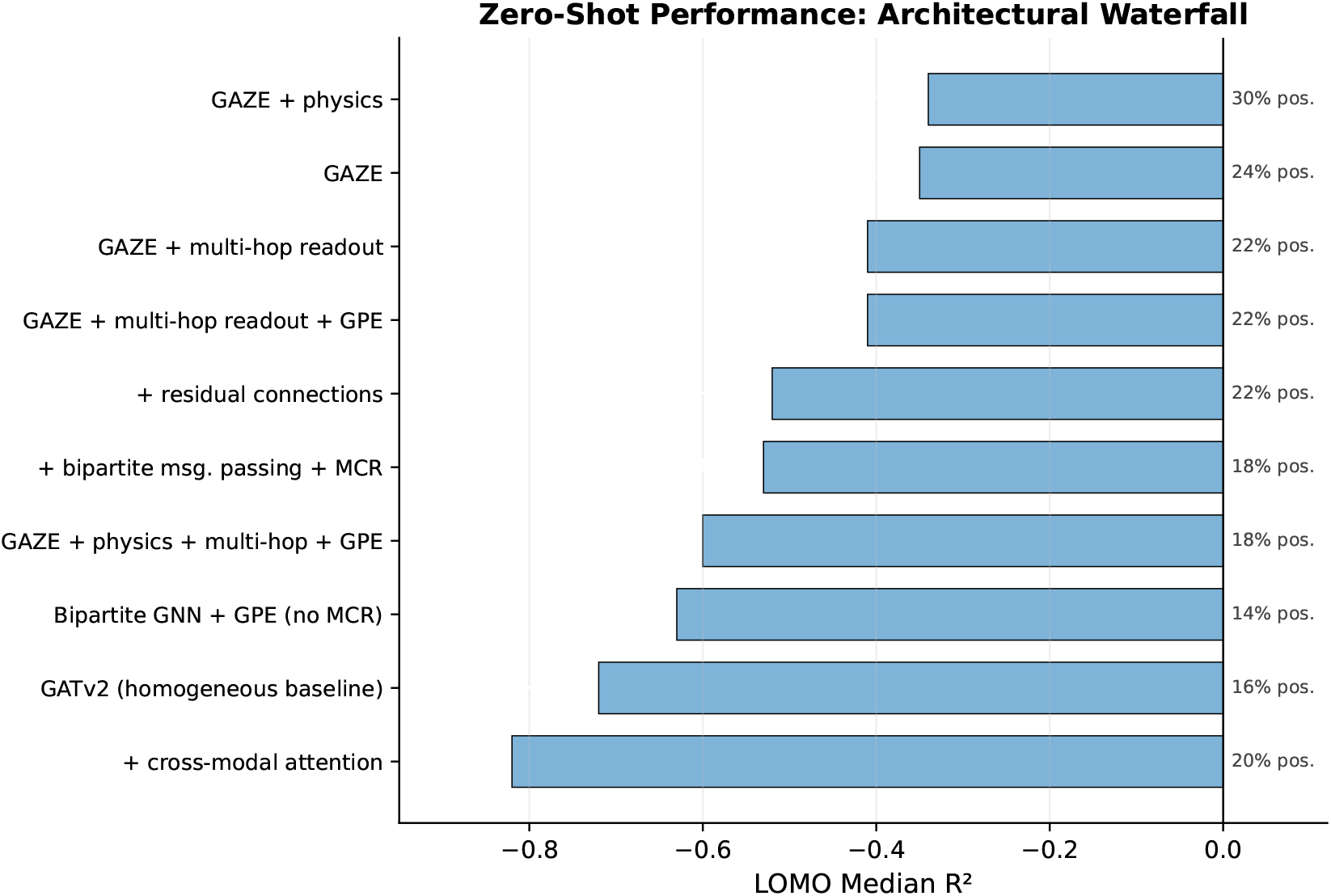
LOMO median R^2^ for all 10 architectural configurations, sorted by performance (best at top). Annotations show the percentage of held-out metabolites with positive R^2^ for each configuration.

GAZE achieved a median Spearman *ρ* = 0.330 across 214 metabolites, with 94% producing positive correlations (**Figure 5A**). The strongest predictions included arachidonate (*ρ* = 0.73), methylimidazoleacetic acid (*ρ* = 0.70), mannose-6-phosphate (*ρ* = 0.70), malonyl-carnitine (*ρ* = 0.69), and glucuronate (*ρ* = 0.67), spanning lipid, carbohydrate, and amino acid metabolism. Performance was consistent across both sub-cohorts (RC18: median *ρ* = 0.34, *n* = 230 metabolites; RC20: median *ρ* = 0.33, *n* = 214). Metabolites that achieved high Spearman *ρ* under LOMO on CAMP also performed well on the external ccRCC cohort (**Figure 5B**).

To place these results in context, we applied all three comparison methods to the same ccRCC tissue data (**Figure 5A**). MEBOCOST achieved a median *ρ* = 0.035 (*n* = 188 metabolites, 58% positive), and scCellFie achieved a median *ρ* = 0.140 (*n* = 123, 71% positive). UnitedMet, evaluated on the same RC18+RC20 data under the same metabolite-holdout protocol used for the CAMP comparison (10-fold CV, *K* = 30), achieved a median *ρ* = −0.017 (*n* = 532, 48% positive), indistinguishable from chance. As on CAMP, this result is consistent with UnitedMet being applied to a metabolite-holdout task for which its latent factorization has no reconstruction mechanism. GAZE, trained on CAMP cell lines without any tissue-specific signal, produced the highest median *ρ* among all four methods on the ccRCC cohort (**Figure 5C**).

## 4 Discussion

GAZE is a physics-informed graph neural network that predicts metabolite concentrations from gene expression, including for metabolites absent from the training set. Existing supervised approaches such as UnitedMet [5] and LASSO regression [4] can impute metabolites present in their training set with high accuracy but operate under a closed-world assumption: they require the target metabolite to have been observed during training. Proxy-based tools like scCellFie [7] and MEBOCOST [8] estimate metabolite activity from enzyme expression without training but correlate weakly with measured concentrations. GAZE addresses the gap between these paradigms by combining transferable chemical representations with physics-informed graph learning, enabling zero-shot prediction of metabolites the model has never seen.

We showed that graph topology and gene expression are the dominant predictive signals, while EC2Vec and SMILES embeddings serve a different function: they are the mechanism through which the model generalizes to unseen metabolites. The zero-shot components (cross-modal attention, MCR, SMILES masking) do not compromise in-distribution accuracy, achieving R^2^ = 0.816 compared to 0.835 for the per-metabolite XGBoost baseline (*p* = 0.35). Although zero-shot metabolite prediction remains difficult and most held-out metabolites produce negative R^2^, our calibration analysis revealed that this largely reflects a shift in mean and scale rather than a failure to learn meaningful patterns: 77% of metabolites with negative R^2^ exhibited significant positive correlation with the true concentrations. The model captures biologically meaningful covariation but cannot anchor predictions to the correct absolute concentration range without having observed any training examples for that compound.

Our mechanism analysis identified a hierarchy of prerequisites for zero-shot success: a valid SMILES embedding is necessary for the MCR to function, the target metabolite must exhibit sufficient biological variability across samples, and smaller molecules are predicted more accurately. Chemical similarity to the training metabolome, graph degree, and enzyme signal strength showed no significant relationship with predictive success. The consistently successful metabolites all occupy fixed positions in linear or near-linear pathway segments at moderate graph degree, where flanking metabolites that remain in training constrain the held-out node through stoichiometric relationships. This indicates that the model generalizes primarily through local pathway interpolation rather than globally transferable gene-to-metabolite rules. Conversely, metabolites without resolved SMILES and large, structurally complex molecules represent systematic failure modes that define the current scope of the approach, whereas highly connected hub metabolites instead plateau at near-zero R^2^, a ceiling rather than a catastrophic failure.

The external validation on ccRCC tissue demonstrated that representations learned from cancer cell lines transfer to an independent biological domain without fine-tuning. On this cohort, GAZE outperformed UnitedMet even though UnitedMet was trained and evaluated on the same ccRCC data. This result is consistent with the zero-shot learning literature [6]: without side information describing the target metabolite, models that rely solely on co-occurrence statistics cannot generalize to unseen items.

Limitations have to be considered. The CAMP dataset comprises only 867 cancer cell lines and approximately 180 metabolites per sample, constraining both sample diversity and chemical coverage; larger multi-tissue datasets may improve generalization. The LOMO protocol tests generalization to metabolites already present in the graph but excluded from training; generalization to metabolites entirely absent from the metabolic reconstruction would require additional graph expansion strategies. The physics loss terms operate on predicted flux values whose ground truth is unavailable, relying on indirect supervision through mass balance and gene–flux correlation constraints. Extending to single-cell and spatial transcriptomics, where dropout noise, cell-type heterogeneity, and domain shifts in gene expression distributions introduce further variance, remains an open direction.

## Data and Code Availability

GAZE is available as a pip-installable Python package with pretrained weights at https://github.com/nrclaudio/ GAZE. The complete training, evaluation, and analysis code to reproduce all results is available at https://github.com/nrclaudio/gaze-reproducibility. The CAMP dataset is publicly available from Benedetti et al. [12]. The UnitedMet ccRCC cohort is available from Xie et al. [5].

## Author Contributions

C.N.-R.: Conceptualization, methodology, software, investigation, formal analysis, data curation, writing—original draft, and writing—review and editing. T.R.: Writing—review and editing. A.M.: Conceptualization, writing—review and editing, supervision, and project administration.

## Acknowledgements

This work was performed using the compute resources from the Academic Leiden Interdisciplinary Cluster Environment (ALICE) provided by Leiden University. The Novo Nordisk Foundation Center for Stem Cell Medicine (reNEW) is supported by Novo Nordisk Foundation grants (NNF21CC0073729). T.R. is supported by the European Union through an ERC grant (SPARK 101140863). Views and opinions expressed are however those of the author(s) only and do not necessarily reflect those of the European Union or the European Research Council Executive Agency. Neither the European Union nor the granting authority can be held responsible for them.

## A Supplementary Figures

